# The homeobox transcription factor HbxB coordinates distinct gene regulatory networks for asexual development and secondary metabolism in *Aspergillus nidulans*

**DOI:** 10.1101/2025.09.26.678847

**Authors:** Emmanouil Bastakis, Rebekka Harting, Tanja Lienard, Merle Aden, Verena Grosse, Blagovesta Popova, Gerhard H. Braus

**Affiliations:** Department of Molecular Microbiology and Genetics, Institute for Microbiology and Genetics, University of Göttingen, Göttingen, Germany

**Keywords:** Fungal transcription factors, asexual development, *Aspergillus nidulans*, *hxbB*, *sclB*, *msnA*, *vapA*, *flbA*, *flbC*, *ppoC*

## Abstract

Formation of conidia as asexual spores and sometimes worldwide distribution through the air is a very important feature of the fungal life style. This process is controlled by several regulatory proteins, including homeobox domain transcription factors. HbxB is one such regulator with implications in the control of development, secondary metabolism and various stress responses in the filamentous fungus *Aspergillus nidulans*. However, the molecular mechanism of the regulatory role of HbxB during asexual development is still elusive. Here we show that HbxB is a nuclear localized protein with great impact on asexual sporogenesis. Employment of high throughput assays like chromatin immunoprecipitation (ChIP-seq) and transcriptomics (RNA-seq), elucidated the *in vivo* binding landscape of HbxB in a genome-wide scale. A set of 238 genes as direct targets of HbxB were identified. A nine bases DNA motif where HbxB prefers to bind *in vivo* was discovered as HbxB response element (HRE). HbxB is influencing the expression of genes encoding master regulators of the asexual development such as SclB, PpoC, FlbA and FlbC. Moreover, the direct transcriptional control of the secondary metabolites sterigmatocystin and emericellamides biosynthesis by HbxB was discovered. Lastly also a previously elusive mutual regulatory control circuit between HbxB and two major regulators of the asexual development SclB and MsnA was found. Both of these regulators can directly induce the expression of *hbxB*. This study provides a detailed molecular mechanism on how HbxB controls *A. nidulans* asexual sporulation.

**Importance:** Fungal distribution mainly relies on the formation of spores that are subsequently dispersed in different media to ensure colonization of substrates and the survival of the fungus. The asexual developmental program is a widely used strategy in the fungal kingdom for production of spores (conidia). The HbxB transcription factor is a nuclear localized, <u>h</u>omeo<u>b</u>o<u>x</u> domain protein, with a strong impact on asexual sporulation of presumably numerous fungal species. This study enhances our understanding of the mechanism with which HbxB exerts its regulatory actions. HbxB is binding *in vivo* to specific DNA regulatory elements of genes encoding proteins with key roles in asexual development (like SclB, MsnA and PpoC), secondary metabolism (such as genes from the sterigmatocystin and emericellamide clusters) and stress response/tolerance. Overall, these findings open a window into how Hbx regulators orchestrate and coordinate fungal asexual developmental programs genome-wide at the molecular level.

## Introduction

Fungi are sessile eukaryotic organisms, which mostly use spores for propagation and to conquer new environments. Based on internal or external stimuli, fungi need to be agile in perceiving and coordinating external signals such as time, medium and environmental conditions for responses like germination or precautions for survival (1). This agility often relies on the choice to increase or decrease the ratio between r distinct developmental programs that will facilitate propagation under challenging conditions.

The asexual program is frequently used for propagation in most fungi, resulting in spores (conidia), which are easily and quickly dispersed through air or water (2). In the filamentous fungus *A. nidulans*, asexual development has been extensively studied. The central regulator is the BrlA (bristle A) transcription factor (3). There are also upstream developmental activators of BrlA (3–6). Among their roles is to ensure that expression of *brlA* is induced under asexual conditions. These regulators are also inhibiting vegetative growth. Known representatives of this group are the Flbs (fluffy low brlA) FlbB, FlbC and FlbD activator proteins (5, 6). Another group of regulators whose transcription is controlled by BrlA and which are crucial for the formation of the asexual structures, alongside with the maturation of the produced spores, are belonging to the so-called central developmental program (3). Apart from BrlA other master regulators of the asexual developmental program are AbaA (abacus A) (7) and WetA (wet-white A) (8). BrlA, AbaA and WetA have the ability to differentially and mutually transcriptionally cross regulate each other to orchestrate and finalize the asexual program (3, 8).

Except of the upstream developmental activators and members of the central developmental program there are also more transcription factors that can greatly influence A. *nidulans* asexual development. The zinc finger transcription factor MsnA has a great impact on the direct *in vivo* regulation of genes encoding major regulators of the asexual program on a genome-wide scale (9). MsnA can bind to the promoter of genes such as *brlA*, *wetA* and *flbs in vivo* under asexual growth conditions and change their expression. SclB (sclerotia like B) is another important transcription regulator of asexual developmental for which the *in vivo* binding landscape during the transition from vegetative to asexual development was elucidated (10). Among other genes that were found to be controlled directly by SclB, *brlA* and *ppoC* (psi factor producing oxygenase C) were some of its top targets. PpoC is a dioxygenase responsible for the synthesis of the precocious sexual inducer B (psiBβ), a secondary metabolite, which promotes asexual conidiation (11, 12).

The homeobox domain (HbxB) transcriptional regulators are present to the majority of multicellular organisms (13, 14). Their function is tightly linked to a broad spectrum of the lifès phenomena such as differentiation, development, stress and others. They carry a homedomain of approximately. 60 aa, which has a helix-turn-helix motif via which it can associate to DNA (14). Hbx proteins have been studied in several fungal species. In *S. cerevisiae* for example they have been linked to mating and growth (15). In *A. flavus* they have been associated to sclerotium and conidiophore formation and with the synthesis of aflatoxin (16). In *A. fumigatus*, they have roles in development, virulence and secondary metabolism (17). In *A. nidulans* eight Hbx proteins have been identified and characterized (18). They include HbxA and HbxB, which show promote the production of the aflatoxin family compound sterigmatocystin, stress response as well as asexual and asexual developmental programs (18, 19).

This study uncovers, a direct *in vivo* regulatory mechanism through which HbxB exerts changes in the expression of particular genes related with *A. nidulans* asexual development, secondary metabolism and stress response. A new transcriptional regulatory circuit among HbxB, SclB and MsnA transcription factors was discovered. This circuit potentially works as an orchestration center where part of the asexual developmental coordination is taking place. A direct implication of HbxB into the synthesis of sterigmatocystin and emericellamides combined with a direct regulation of stress-related genes was discovered.

## Results

### The HbxB transcription factor preserves its nuclear localization during vegetative and the asexual growth

The roles of homeobox transcription factors in fungi are tightly connected with development (20). Previous genetic studies revealed the impact of HbxA (AN1217) and HbxB (AN2020) on the development and the secondary metabolism of *A. nidulans* (18). Here we set to understand the mechanism through which, specifically HbxB is able to control asexual development and secondary metabolism of the fungus. The open reading frame of the *hbxB* gene gives an 1824 bp transcript, consisting of three exonic and two intronic regions (**FIG 1A**). This transcript, when translated, leads to the synthesis of a 293 aa protein, which is constituted by the homeobox domain, as predicted by the ScanProsite algorithm of the Prosite data base (21) and a nuclear export signal as predicted by the LocNES algorithm (22), respectively. Freely available web tools did not predict a nuclear localization signal (NLS) in HbxB. It was hypothesized that HbxB functions might be controlled by posttranslational modifications. In-silico prediction analysis for putative phosphosites in the HbxB protein was conducted. The NetPhos tool (23) suggested at least four serine residues of HbxB which might be phosphorylated by kinases (**FIG 1A**). It has been demonstrated previously that the *hbxB* deletion strain shows severe defects, particularly in the production of conidia when it is growing for 5-d under asexual conditions (19). For this study, the corresponding phenotype of the deletion strain was studied at earlier and later time points during asexual growth. Plates were spot-inoculated and grown for 3- and 8-d under asexual conditions, correspondingly. The observed phenotype of the *hbxB* deletion strain compared wildtype after 5-d of incubation was similar to earlier studies (Son et al., 2021). The deletion strain had a fluffy appearance and the number of conidia produced was dramatically reduced compared to the wildtype for both time points examined (**FIG 1B-E**). The *hbxB* deletion strain was transformed in locus with a transgene coding for HbxB-GFP, which was used as complementation strain. The phenotype of this strain fully complemented the severe phenotype of Δ*hbxB*, supporting that HbxB was responsible for the observed phenotype of the deletion strain and that the GFP-fusion is functional (**FIG 1B-E**).

**FIG 1:**
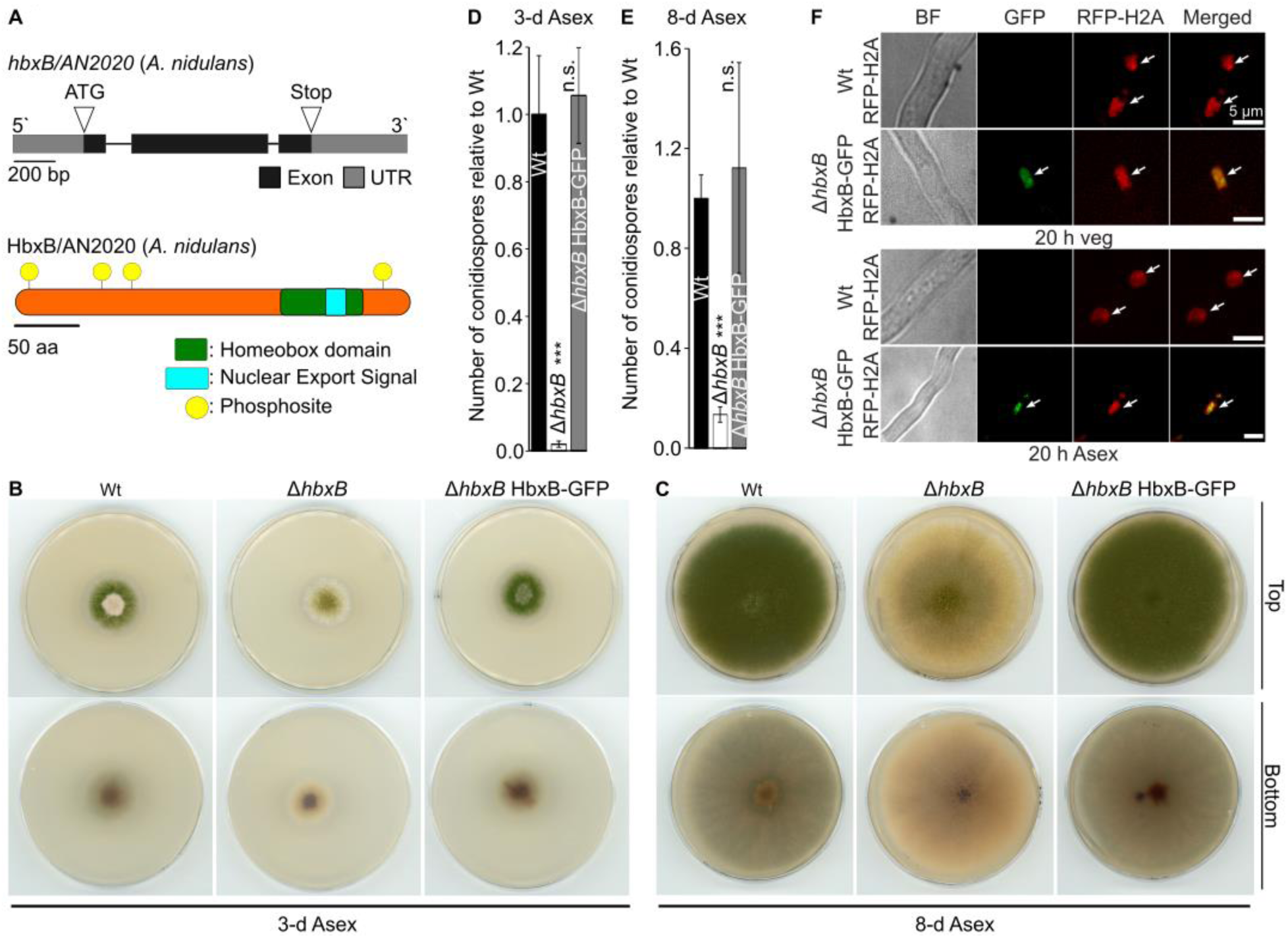
HbxB is a nuclear localized protein involved in asexual conidiation of *A. nidulans*. (**A**) Representation of the open reading frame of *hbxB* (*AN2020*) and the composition of predicted domains, nuclear export signal and putative phosphosites of the encoded protein HbxB. Phenotypical characterization of the Δ*hbxB* deletion and the Δ*hbxB* HbxB-GFP complementation strain after 3-d (**B**) and 8-d (**C**) of asexual growth with light and at 37 °C. For both groups initially 2000 spores were spotted in the middle of the plates and scans photos were made from the top and the bottom of each plate, after the indicated growth periods. Quantification of the total number of conidia relative to Wt (wildtype), produced by the deletion (Δ*hbxB)* and the complementation strain (Δ*hbxB* HbxB-GFP), grown for 3-d (**D**) and 8-d (**E**) under asexual growth conditions. Initially 30,000 spores for each strain spread homogenously all over the plates. The number of conidiospores per plates was quantified after the indicative number of days. A pairwise comparison among of the deletion and complementation strains with the Wt was performed by student t-test: ***: p-value<0.001 and n.s.: not significant. (**E**) Images derived from live imaging confocal microscopy show the nuclear localization of the HbxB transcription factor of vegetatively (upper side) and asexually (bottom side) grown hyphae. Initially, spores were inoculated in liquid medium and grown for 20 h under vegetative and for 20 h under asexual development inducing conditions. Signals from the green channel illustrate GFP and signals from the red channel illustrate the common nuclear marker H2A fused to RFP. White lines present a length of 5 µm. White arrows point to the nuclear localized HbxB-GFP protein and the nuclear marker RFP-H2A.

The nuclear localization of HbxB was addressed by ectopically co-expressing a transgene for the global nuclear marker of H2A (Histone 2A) fused with the red fluorescent protein (RFP) to the Ń-terminus in the complementation strain HbxB-GFP. Hyphae from that strain and a Wt strain also expressing RFP-H2A were examined by confocal live imaging microscopy after 20 h of vegetative growth and after 20 h of asexual growth. In both cases HbxB had a nuclear localization (**FIG 1F**).

In conclusion, the role of HbxB is strongly associated with the conidiation of the fungus in earlier and later points than the one of 5-d studied before. Moreover, it was shown for the first time that HbxB is nuclear localized not only during vegetative but also during the asexual growth of *A, nidulans*.

### A group of 238 genes are direct *in vivo* targets of HbxB during asexual development of *A. nidulans*

Apart from the characterization of several homeobox transcription factors in fungi and specifically in *A. nidulans*, to our knowledge there is a lack of studies showing the *in vivo* binding landscape of such regulators in a genome-wide scale. To this end it was decided to generate the binding profile of the HbxB transcription factor at an early step of the asexual growth by an *in vivo* ChIP-seq assay. The ChIP-seq experiment with the complementation strain of HbxB (Δ*hbxB* HbxB-GFP) was performed with mycelia grown for 9 h asexually, to discover the *in vivo* target genes of the HbxB transcription factor. For the ChIP experiment four independent biological replicates of HbxB-GFP (anti-GFP antibody applied during the IP) with the corresponding inputs (anti-GFP antibody not applied) were used to subtract any background/noise signal. The analysis was conducted to identify statistically significant peaks (position in the genome where the *in vivo* binding of HbxB occurs) in all four independent sets. A total number of 1115 of locus gene IDs, direct targets of HbxB, appeared reproducibly in all sets (**FIG 2A**). All the identified peaks where located into regions spanning up to 3 kb from the transcription start site (hear after TSS) of the genes. Distribution analysis of all the identified peaks over different genomic elements highlighted that around 70 % of them is located to promoter regions up to 1 kb from the TSS (**FIG 2B**).

The DNA motif with which HbxB prefers to be associated to *in vivo* is still elusive. As part of the ChIP-seq analysis, 150 sequences of 100 bp each were extracted, from the 150 top-scored peaks of each independent set of analysis. All these sequences were located below the summit of the corresponding identified top peaks. In fact, each of the 100 bp sequences (from all sets), was defied as a fragment that spans 50 bp downstream and 50 bp upstream from the exact position of the corresponding summit of each peak. These sequence groups were used as inputs for performing a *de novo* motif discovery by the RSAT webtool (24). The motif that emerged consistently in all independent sets had a consensus sequence of nine bp: 5’-AATTCCCCG-3’; we named this identified DNA motif as HbxB response element (HRE) (**FIG 2C and FIG S1A**).

*In vivo* binding of a TF to a regulatory region of a gene, does not necessarily mean differential expression of the target. To test which of the 1115 genes, where HbxB was found to be associated with the promoters (**FIG 2A**), is differentially expressed, we performed RNA-seq experiments under the same conditions as the ChIP-seq with HbxB-GFP. RNAs from Wt and Δ*hbxB* strains were extracted from mycelia growing for 9 h under asexual conditions. A principal component analysis (PCA) with all biological replicates of both strains was performed to test the integrity of the samples from the RNA-seq experiments. This analysis indicated that three biological replicates from the Wt samples and four biological replicates from the Δ*hbxB* samples were grouped into two distinct separate clusters, confirming the validity of the whole experiment (**FIG S2A**). A subsequent step of the analysis has shown that 2361 genes are getting differentially expressed (applied cut offs: p<0.05 and −1 ≤ log_2_FC(Fold Change) ≥ 1) when comparing the Δ*hbxB* with the Wt samples (**FIG S2B**). From the total number of differentially expressed genes (hereafter DEGs), 1437 were found to be upregulated and 924 downregulated. A gene ontology (GO)-enrichment analysis was conducted by employing the ShinyGO algorithm to examine what type of proteins are encoded by those 2361 DEGs (25). The majority of the statistically significant overrepresented categories of the biological processes (BP), were related with steps and pathways of fungal secondary metabolism (**FIG S2C**).

An overlap between the 1115 target genes found in ChIP-seq with the 2361 DEGs from the RNA-seq was performed to identify the genes with HbxB *in vivo* binding at the promoters that resulted in a change in expression. A group of 238 genes emerged from this overlap (**FIG 2D and 2E**). A heatmap was generated based on the expression of those 238 genes. It highlighted the strong and distinct expression profile of the Δ*hbxB* versus that of the Wt samples (**FIG 2E**).

In sum, the HbxB transcription factor can indirectly and directly influence the expression of many genes in the genome of *A. nidulans* during asexual growth. The great majority of these genes are associated with processes related with the secondary metabolism of the fungus.

### The asexual developmental program of *A. nidulans* is transcriptionally controlled by the HbxB transcription factor

The main focus of this study was to understand the mechanism through which the HbxB regulator is particularly controlling *A. nidulans* asexual development. Hence, a search in data sets of the ChIP-seq and RNA-seq for genes was conducted that particularly encoding known asexual regulators. HbxB seems to directly control the expression of several known asexual regulators *in vivo* through promoter binding (**FIG 3A and 3B**). In fact, its binding to the promoter regions of *sclB*, *flbC* and *ppoC* causes induction of these genes. A similar *in vivo* association with the promoters of *flbA* and *vapA* can cause the repression of these two genes, correspondingly. Further analysis has indicated that there were also genes encoding other asexual regulators (*rodA*, *dewA*, *yA*, *sfgA*, *abaA*, *ppoB* and *wetA*), where the expression was found to be altered in our RNA-seq data (**FIG 3B)**, although, there was no *in vivo* association of HbxB with their promoter regions (**S3 Table)**. Moreover, there were also several genes found that encode established asexual regulators where HbxB can bind *in vivo* to their promoter regions. However, this binding did not lead to differentially expression; at least not at the time point of our RNA-seq analyses (**FIG 3C and S4 Table**). Overall these results show that at relatively early steps during *A. nidulans* asexual growth, the HbxB transcription factor can directly or indirectly affect the expression of genes encoding regulatory proteins of the asexual developmental program.

### HbxB is promoting the synthesis of meroterpenoids, sterigmatocystin and emericellamides

The fungal secondary metabolism plays an important role in the communication of fungi with other organisms and ultimately for their survival. Molecules that are synthesized by the secondary metabolism influence development, the defense of the fungus and the interplay with other living organisms (12, 26). The ChIP-seq data were examined for binding events of HbxB to promoters of genes coding for members of known secondary metabolite gene clusters to estimate the impact of HbxB on secondary metabolism during asexual growth.

It was found that the sterigmatocystin (*ste*) and the emericellamide (*eas*) gene clusters accommodated multiple binding events of HbxB, to regulatory regions of several members (**FIG 4A and 4B**). It was then tested, if these *in vivo* binding events, lead to changes in the expression of genes located nearby. The transcriptomic profile of the Δ*hbxB* versus Wt, during asexual development has shown that all the members of the *eas* cluster were downregulated, meaning that the role of HbxB is to induce their expression during asexual growth (**FIG 4C**). In the case of the sterigmatocystin gene cluster, there were a plethora of binding events by the HbxB, across the whole cluster, but only four gene members (*AN7822*, *aflR AN11021*, *stcA*) were found to be repressed in the RNA-seq data (**FIG 4C**). Noteworthy to mention here, among these four genes *stcA* is coding for the backbone enzyme of the whole cluster (27, 28) and *aflR* codes for the main transcriptional regulators of the cluster (29). Additionally, HbxB was found to be associated with the *stcA* and *aflR* promoters, implying that the differential regulation of the corresponding genes relied on this direct *in vivo* binding events (**FIG 4A**). All four independent sets of the ChIP-seq analysis showed binding of HbxB to the *stcA* promoter. In the case of the *aflR* promoter region three out of the four sets of the ChIP-seq analysis showed a statistically significant binding of HbxB (**FIG S4 and S3 Table**).

This influence of HbxB on the *ste* and *eas* clusters on a molecular level might have a functional impact for the fungus. Secondary metabolites from Wt, Δ*hbxB* and Δ*hbxB* HbxB-GFP strains grown for 3-d under asexual growth conditions were extracted and subsequently analyzed to test this hypothesis. LC-MS/MS analysis revealed that sterigmatocystin was the metabolite found to be affected the most in the *hbxB* deletion strain compared to Wt. Emericellamides C/D and E/F were also found to be significantly reduced (**FIG 4D**). The secondary metabolite analysis showed that the meroterpenoids austinol and dehydroaustinol were significantly decreased as well in the Δ*hbxB* strain compared to the levels that the Wt showed. However, no binding of HbxB to any of the promoter regions of genes from the *aus* cluster has been identified. But many of the *aus* cluster genes were found to be strongly differentially expressed in the corresponding transcriptomic data (**FIG 4C**).

To sum up these results, HbxB influences the expression of genes of several secondary metabolite gene clusters and this influence has a direct impact on the synthesis of the corresponding metabolites during fungal asexual development.

### A molecular circuit between the HbxB, SclB and MsnA transcription factors coordinates the asexual developmental program of *A. nidulans*

The *in vivo* binding landscapes in the genome of *A. nidulans* of two major asexual regulators, SclB and MsnA, were recently uncovered (9, 10). It was tested whether SclB and MsnA might be able to control the expression of *hbxB* as regulatory gene of the asexual developmental program. An analysis with our ChIP-seq data of the SclB (PRJNA1292721) and MsnA (PRJNA1198984) transcription factors, deposited at the sequence read archive of NCBI was performed. It was found that SclB is strongly associated with the promoter of *hbxB* during vegetative growth and early asexual development (**FIG 5A**). To further test if these binding leads to a transcriptional response, our transcriptomic RNA-seq data (also deposited at NCBI under the ID: PRJNA1293339) from the same study was analyzed. The expression of *hbxB* was examined under both growth conditions. During vegetative growth *hbxB* was strongly downregulated in the Δ*sclB* strain compared to Wt (**FIG 5A**). However, *hbxB* expression did not change when the RNA-seq data from early asexual growth were examined (data not shown). Analysis of the ChIP-seq data for the MsnA master regulator was conducted to analyze whether *hbxB* was one of its targets. This analysis showed that MsnA binds to the promoter region of *hbxB*, not only during vegetative growth but also during early asexual development of the fungus (**FIG 5B**). To examine if these binding occurrences have transcriptional impact on the expression of *hbxB*, RNAs were extracted from mycelia of Wt and Δ*msnA* strains, growing either under vegetative or asexual conditions correspondingly and a qRT-PCR analysis was performed. The expression of *hbxB* was found to be strongly induced in the *ΔmsnA* strain compared to Wt, when vegetative mycelia were tested (**FIG 5C**). However, the *hbxB* expression was not changed when mycelia from the same strains were tested under asexual growth conditions (**FIG 5C**). Next, it was questioned whether HbxB could influence the expression of *msnA* as well. A search in the NGS data generated for this study showed that HbxB can be associated to multiple positions of the *msnA* promoter region under asexual growth (**FIG 5D**). Additionally, when the corresponding transcriptomic data from the identical growth conditions were examined, *msnA* was not found to be differentially expressed, at least under the applied cut offs: p < 0.05 and −1 ≤ log_2_FC(Fold Change) ≥ 1. Lastly, it was discovered that HbxB is strongly binding to its own promoter (**FIG 5D**), a fact that would imply an autoregulatory role that HbxB might exert towards its own expression. Summarizing, these results propose a new, so far elusive trimeric transcriptional regulatory network, among the transcription factors HbxB, SclB and MsnA, which coordinates *A. nidulans* asexual development, early on from vegetative growth.

### Genes related to several stress responses and stress tolerance are differentially expressed by HbxB during *A*. *nidulans* asexual growth conditions

A former study has examined the implication of the HbxB regulator in stress responses of *A. nidulans*. It was found that HbxB has a role in the tolerance of thermal and oxidative stress (18). A subsequent study has partially focused on the transcriptional responses mediated by HbxB in *A. nidulans* spores, with emphasis on various stress-related genes and pathways (19). The material from which we generated our ChIP-seq and RNA-seq data was asexually grown hyphae and not spores. However, it was hypothesized that perhaps HbxB is able to occupy promoters of stress-related genes and subsequently change their expression. To test this, transcriptomic levels of 44 genes stress related (**S6 Table**) were examined. These genes are associated to several stress response pathways related to temperature, oxidative stress, tolerance against radiation/UV but also included core genes of the HOG (High Osmolarity Glycerol) signaling, trehalose synthesis and beta-glucan degradation genes. It was found that only nine (*catA*, *catC*, *exgB*, *exgC*, *treB*, *ccg9*, *nopA*, *hsp20* and *hsp140*) of these 44 genes were differentially expressed in our transcriptomic analysis performed with RNAs from the Δ*hbxB* and Wt strains grown asexually (**FIG S3A**). The ChIP-seq data of HbxB were investigated to analyze whether the expression of the nine genes is changed because of the presence of HbxB at their promoters. Only the differential expression of *ccg9* was found to be changed due to HbxB binding to its promoter (**FIG S3B**). When a search of the 44 stress-related genes in the list of 1115 reproducible targets of the HbxB from the ChIP-seq data was conducted, it was found that HbxB *in vivo* associates with seven of these genes: *hbxA*, *bglH*, *crhB*, *pbsB*, *ccg9, catB* and *treA* (**FIG S3B**). Only *ccg9* transcription was found to be differentially expressed under the same conditions. In summary, these data revealed that HbxB is able to indirectly affect the expression of several stress-related genes during asexual growth conditions. In addition, HbxB remains associated with promoters of several other stress genes *in vivo*, but cannot influence their expression, at least not for the time point under examination. Most likely, this occurs later and presumably in presence of specific stimuli and other co-factors.

## Discussion

Propagation is the most prominent requirement for all the living organisms, including fungi. Otherwise a species will be extinct. Fungal development is tailored to respond quickly to internal and external cues that will facilitate an efficient propagation and colonization. Asexual development is the most prominent fungal developmental program (30). The spores produced are rapidly transported through the air to long distances, helping the fungus to colonize new environments. The regulation of asexual development is controlled and coordinated on multiple levels during the fungal life cycle. In *A. nidulans* asexual development has been extensively studied and several central players have been elucidated. Homeobox (Hbx) transcriptional regulators, comprise a relatively small family of eight member in *A. nidulans*. Although, the majority of them have been mostly genetically characterized (18), the mechanistic overview of their contribution to development in a genome-wide scale is missing. In this study we show the global impact of the HbxB transcription factor, in the regulation of genes during asexual development and secondary metabolism of the fungus.

To our knowledge this is the first study, that proves a clear nuclear localization of HbxB protein from *A. nidulans*. HbxB is localized to the nucleus prior to the initiation of asexual developmental, when grown vegetatively (**FIG 1**). This implies that its role in *A. nidulans* development in starts before the fungus makes the final decision which program to follow. In a previous study, where the expression of *hbxB* was investigated in mycelia grown under asexual conditions, mRNA levels of *hbxB* were increasing even up to 24 h post asexual induction (18). Approximately 24 h is the overall time that is needed for the completion of the asexual developmental program, If the mycelium is developmentally competent before (2, 12). The results from our study together with the findings by Son et al, highlight the role of HbxB in asexual growth spanning from its onset till its completion. The direct implication of HbxB in the final steps of asexual development, it constitutes a worthy subject for future investigation.

Studies investigating the genome-wide *in vivo* binding profiling of Hbx proteins in fungi are very restricted. Via a ChIP-seq assay regulatory binding positions of an Hbx protein from the plant pathogen *Fusarium graminearum* to promoters of 142 differentially expressed genes were identified (31). To our knowledge similar studies of any of the eight HbxB proteins were not carried out. Therefore, we generated a binding profile for the *A. nidulans* HbxB in the context of asexual development. The ChIP-seq data of this study showed that at the relatively early steps of the asexual growth, HbxB is reproducibly and *in vivo* associated with promoters of 1115 genes (**FIG 2**). To conclude, which of these binding events leads to changes in the expression of genes nearby, corresponding transcriptomic (RNA-seq) data were also generated (**FIG S2**). The combination of ChIP-seq and RNA-seq data has revealed a set of 238 genes, where HbxB binds to the promoters and this association triggers immediate changes to the expression (**FIG 2**). Fan et al., found in *F. graminearum* a similar relatively small set of genes (142) where the association of Htf1(Homeobox transcription factor 1) caused direct changes to the expression (31). Our study has also elucidated the, HbxB response element (HRE), a six bp DNA motif, where HbxB is mostly preferring to be associated with in the genome of *A. nidulans* (**FIG 2**). Comparing the HRE of HbxB from *A. nidulans* with the Htf1 response element from *F. graminearum*, it seems that the latter shows preference for an *in vivo* association to DNA sequences which are smaller, 4-5 bp long and with different composition from HRE of *A. nidulans* (31). The GO-enrichment analysis from the direct targets of HbxB showed a predominant preference for genes related to secondary metabolism of *A. nidulans* (**FIG S2**). Genes coding for proteins implicated to the metabolic processes in general have also been emerged from the GO analysis of Htf1 targets from *F. graminearum* (31).

**FIG 2:**
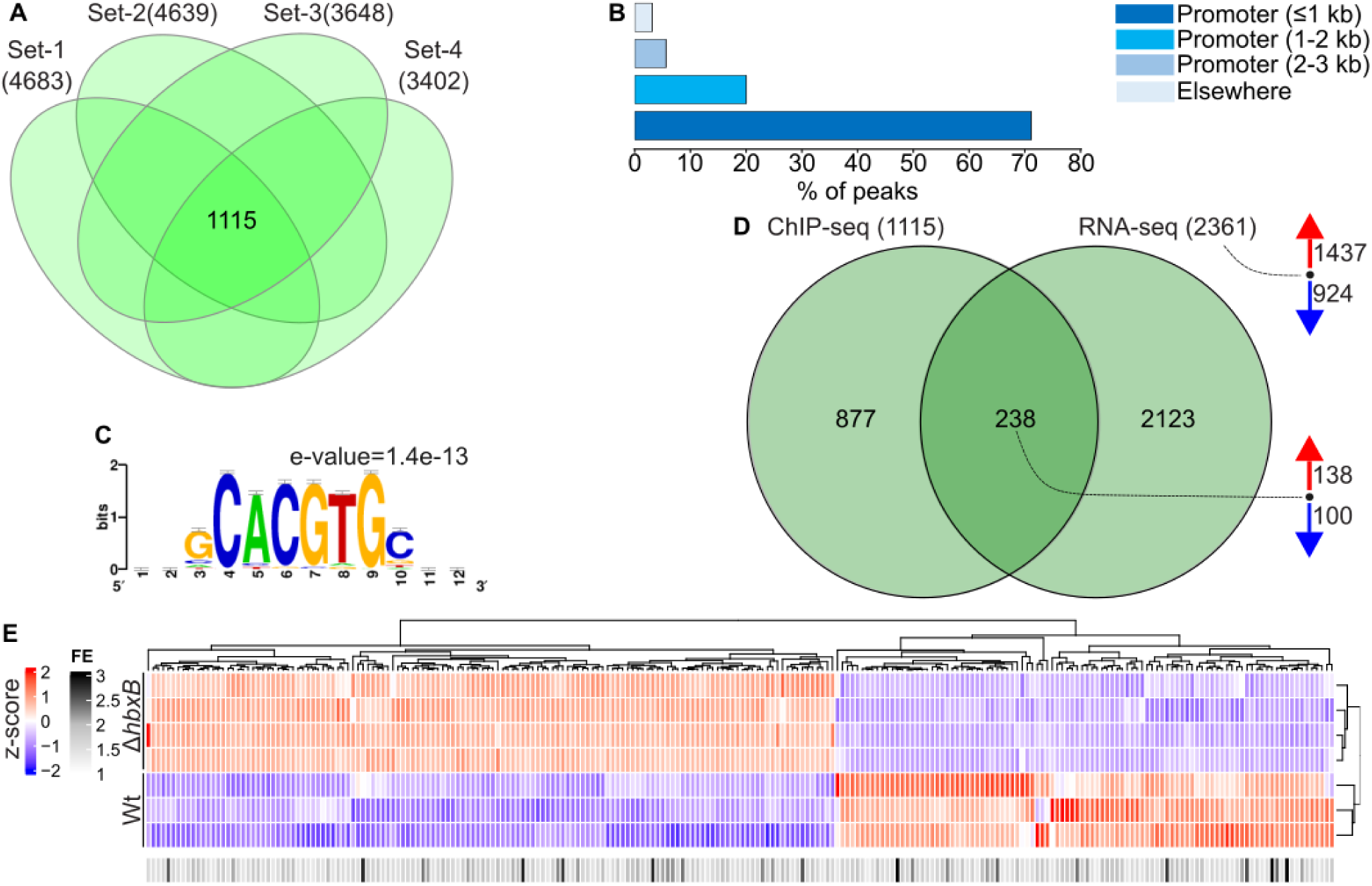
HbxB directly affects, the expression of 238 genes in the *A. nidulans* genome *A. nidulans* during the asexual development. by its *in vivo* binding association to regulatory regions during asexual development. (**A**) The overlap as presented by a Venn diagram, depicts the number genes (locus IDs) found to be targets of HbxB in all four independent sets of ChIP-seq analysis (HbxB-GFP versus input) performed after 9 h of asexual growth. Each set of analysis is composed by genes, where HbxB was found to be associated with their promoter regions up to 3 kb upstream from the TSS (transcription start site) *in vivo*. Moreover, all sets of analysis were primarily subjected to a filtering of the identified peaks (*in vivo* binding positions of HbxB), regarding the p-value<0.05 and the fold enrichment (FE)≥1.2. (**B**) Bar chart presenting the distribution of the identified statistically significant peaks of the ChIP-seq, over different genomic elements of the *A. nidulans* genome. The numbers presented as % derived from the analysis of the representative independent set-1 (out of in total four) of the ChIP-seq data; very similar peak distribution was obtained while performing the corresponding analysis for the remaining three independent sets (FIG S1A). (**C**) Representative motif derived from a *de novo* motif analysis of the set-1 of ChIP-seq data. The motif presents top scored DNA sequence where HbxB was found to be associated with *in vivo*. The analysis was performed with the RSAT webtool, using as input a group of 100 bp sequences extracted under the summit of the first 150 top scored peaks for each independent set of the ChIP-seq analysis. The corresponding *de novo* motif discovery for the remaining three independent sets of the ChIP-seq data is presented in FIG S2B. (**D**) Venn diagram highlights the overlap between the 238 direct target genes of the HbxB transcription factor. As such those are characterized that HbxB was found to be associate to their promoter regions *in vivo* in all four independent sets of the ChIP-seq analysis and at the same time were differentially expressed in the RNA-seq data of Δ*hbxB* versus Wt. Both high throughput experiments were performed with mycelia initially grown for 20 h vegetatively and subsequently transferred to solid medium for another 9 h under asexual development inducing conditions (light and at 37 °C). The applied cut offs for the ChIP-seq were: p<0.05 and FE≥1.2 and for the RNA-seq: p<0.05 and −1 ≤ log_2_FC(Fold Change) ≥ 1. The red arrows at the left side of the Venn refer to the number of upregulated genes and blue arrows to the number of downregulated genes. (**E**) Heatmap with red-blue color gradient shows the gene expression pattern (as z-score values) from the RNA-seq data from **D** of the 238 direct target genes of the HbxB transcriptional regulator. Heatmap with the grey-black gradient presents the binding (as (FE) fold enrichment) of HbxB to the promoters of those 238 genes, as found from the ChIP-seq study in **A**. The fold enrichment was calculated as the average derived from all four independent sets of analysis.

The influence of HbxB on secondary metabolism of *A. nidulans* was examined prior to this study. It was discovered that HbxB is promoting the production of sterigmatocystin during an extended period of 7-d under sexual growth conditions (19). In the current study it was found that HbxB is also able to strongly induce the synthesis of sterigmatocystin under asexual growth conditions (**FIG 4**). Moreover, it was shown that this induction is an immediate consequence of the direct *in vivo* control of HbxB on regulatory regions of genes of the *ste* cluster. These genes encode essential proteins for the synthesis of sterigmatocystin, such as the StcA backbone enzyme and the major transcriptional regulator of the cluster AflR (**FIG 2 & FIG S4**). The implication of HbxB in the regulation of secondary metabolism was further extended, since its promoting role in the synthesis of meroterpenoids, austinol, dehydroaustinol and emericellamides was discovered as well (**FIG 4**). Simultaneous application of dehydroaustinol and diorcinol is able to restore sporulation defects of a strain carrying a gene deletion of the asexual master regulator FluG (32). This further corroborates the direct involvement of HbxB in the asexual development via transcriptional induction of metabolite synthesis that function as inducer of this program. The cyclopeptides emericellamides, are mostly known for their antibiotic activities (33). Because of that, it can be proposed that HbxB broadly strengthens the defense of *A. nidulans* against other organisms by inducing the synthesis of emericellamides, through activation of all four genes of the *eas* cluster (**FIG 4**).

The focus of this study was the disclosure of the mechanism through which HbxB is controlling and promoting *A. nidulans* asexual development. The generated genomic data revealed that there are several genes coding for major regulators of asexual growth, which expression is directly or indirectly changed through HbxB under asexual growth conditions (**FIG 3**). *flb*C, *ppoC* and *sclB* code for known and established players of asexual development. HbxB binds to their promoters, hence activating their expression. However, two additional categories of genes coding for asexual central players were discovered, where HbxB is related to in a different manner. There is a set of genes, such as *brlA*, *vipC*, *flbD* and others, where HbxB binds to their promoter, but there is no change in their expression, at least at the tested time point. This would imply that these *in vivo* binding events become potentially functional, in terms of causing changes in gene expression, at earlier or later time points potentially when other cofactors or stimuli are present. Another set of genes related to asexual development was also noted, where no binding of HbxB was identified to their promoter. Expression of *wetA*, *abaA*, *sfgA* and others was significantly changed in transcriptomic data, suggesting indirect regulation (**FIG 3**). Accumulative these results suggest that the HbxB transcription factor influences asexual growth of *A. nidulans* by altering the expression of key genes in directly but also indirectly manner.

**FIG 3:**
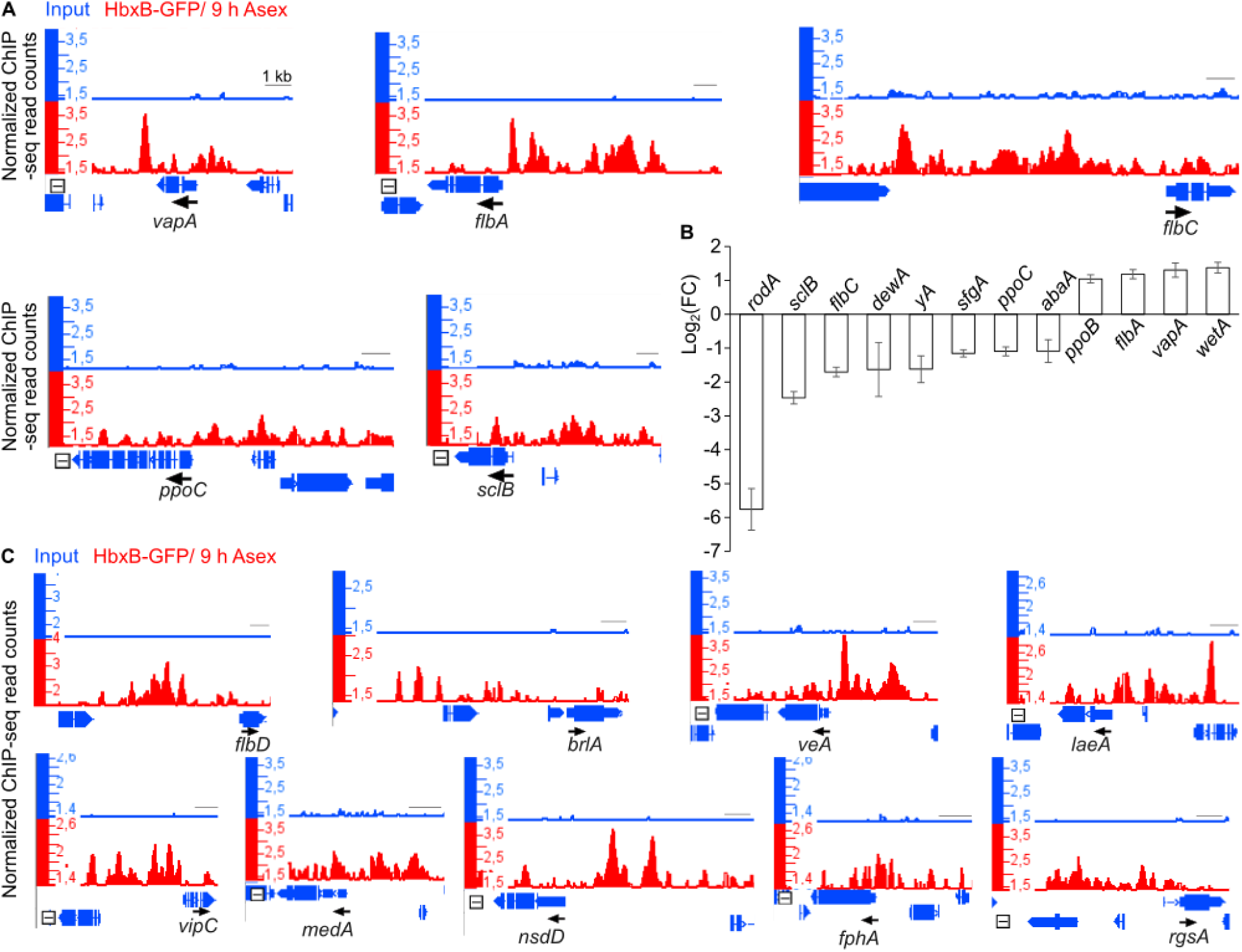
HbxB controls the expression of genes coding for known regulators of asexual development. (**A**) Views from the integrative genome browser (hereafter IGB views) for the ChIP-seq data of HbxB, depicting peaks (*in vivo* binding positions) of the transcription factor to promoter regions of genes coding for known regulators of asexual development such as, VapA, FlbA, FlbC, PpoC, SclB. All these genes are direct targets of HbxB, as presented in **FIG 2E** and **2F**. Blue tracks illustrating the input (background signal) and red tracks the signal derived from the ChIP-seq of the IP with the anti-GFP antibody. Black arrows below the corresponding genes indicate the direction of their transcription. A representative of one set of analysis, out of four in total is presented. The black line, inside each snapshot represents a length of 1 kb in the genome. (**B**) Bar chart showing the expression (as of Log_2_(FC) values) of genes coding for known regulators of the asexual development in *A. nidulans* as derived from the RNA-seq analysis (applied cut offs: p < 0.05 and −1 ≤ log_2_FC(Fold Change) ≥ 1) for mycelia of Δ*hbxB* and Wt strains grown under asexual conditions. All of these genes are transcriptionally regulated by HbxB, some directly (by its *in vivo* binding to their promoters): *vapA*, *flbA*, *flbC*, *ppoC* and *sclB* and others indirectly (no binding to their promoter detected in our ChIP-seq data): *rodA*, *dewA*, *yA*, *abaA*, *ppoB* and *wetA*. (**C**) IGB views, depicting ChIP-seq *in vivo* bindings (peaks) by the HbxB to promoter regions of genes coding for known regulators of asexual development such as, *flbD*, *brlA*, *veA*, *laeA*, *vipC*, *medA*, *nsdD*, *fphA* and *rgsA*. However, these bindings do not lead to immediate changes in the expression of these genes, as it was verified in the corresponding RNA-seq data from the same conditions (**FIG S2** and **S4 Table**).

**FIG 4:**
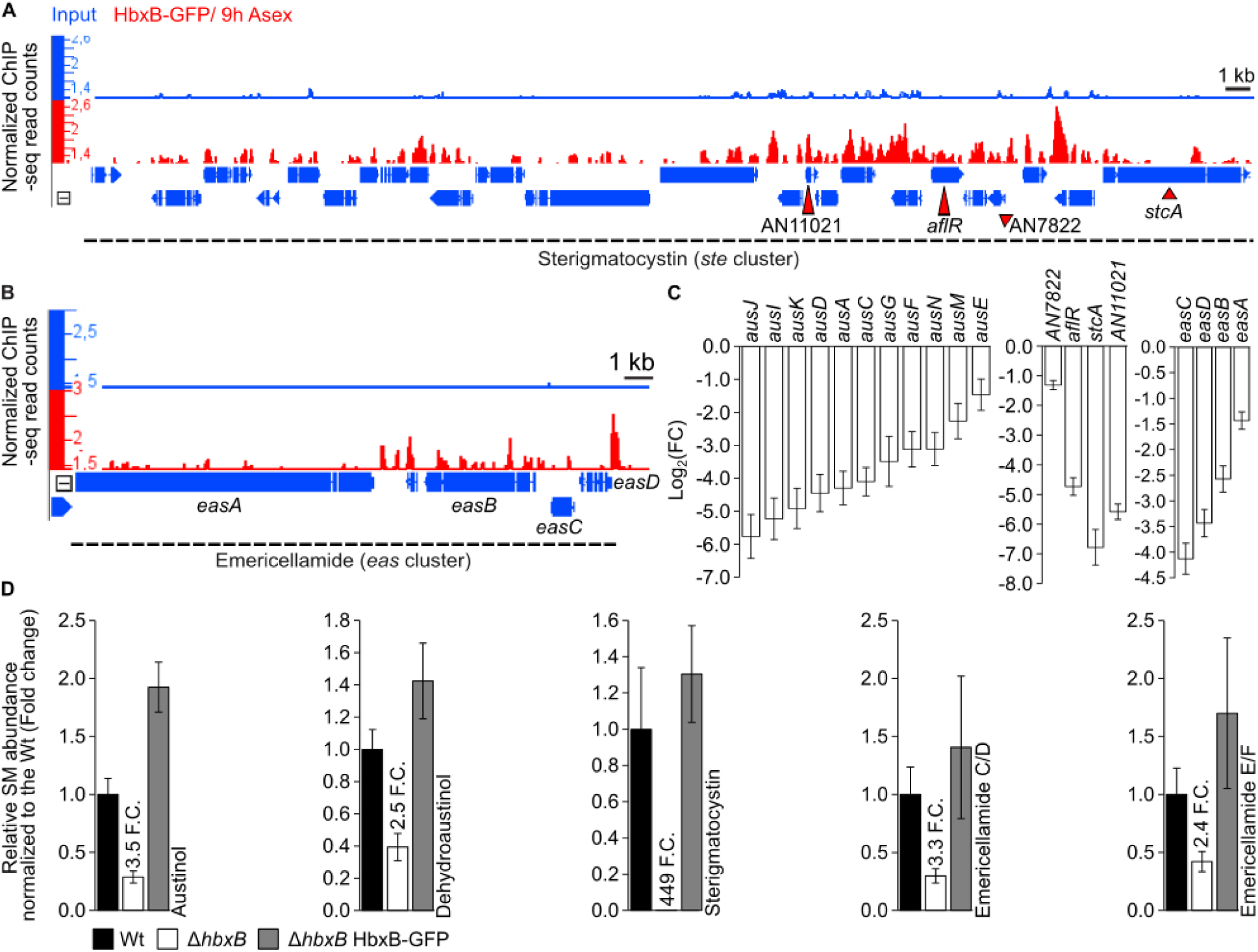
The synthesis of several *A. nidulans* secondary metabolites is transcriptionally controlled by the HbxB transcription factor. IGB views from the IGB depict ChIP-seq data for the *in vivo* association of HbxB to the promoter regions of genes coding for members of the (**A**) sterigmatocystin and (**B**) emericellamide gene clusters. (**C**) Bar charts illustrating the differential expression of genes (as of Log_2_(FC) values), coding for members of the *aus*, *ste* and *eas* gene clusters, from the RNA-seq analysis (applied cut offs: p < 0.05 and −1 ≤ log_2_FC(Fold Change) ≥ 1) performed with RNAs derived from mycelia of Δ*hbxB* and Wt strains, grown under asexual conditions. (**D**) Bar charts showing relative abundance of secondary metabolites expressed as fold change compared to wildtype. The corresponding metabolites were detected by employing a LC-MS/MS platform with a charged aerosol detector (CAD). The extracts of the secondary metabolites derived from the Wt, the Δ*hbxB* and the complementation strain (Δ*hbxB* HbxB-GFP), grown for 3-d under asexual conditions on minimal medium plates. For quantification, four independent biological replicates were used for each strain.

Recently the mechanisms via which the prominent transcriptional regulators of asexual growth, MsnA and SclB are operating in a genome-wide scale were uncovered (9, 10). SclB was found to be a direct target of HbxB for asexual growth. HbxB was also associated to the promoter of *msnA*, without leading to differential expression of the corresponding gene (**FIG 5**). Subsequently, it was hypothesized that HbxB, SclB and MsnA might be main components of a regulatory circuit, which coordinates asexual development in *A. nidulans*. Taking advantage of the ChIP-seq and transcriptomic data of SclB and MsnA, it was possible to reanalyse them to discover a regulatory network among the three transcription factors. In fact, not only SclB but also MsnA can directly control the expression *hbxB*, by *in vivo* binding to its promoter region (**FIG 5**). In that way, this study has connected three main regulators of asexual growth, the homeobox HbxB, the zinc-cluster (C6) SclB and the zinc-finger MsnA transcription factors. An additional finding was that HbxB, like its two partners from the same circuit can potentially transcriptionally be autoregulated, since they can all be associated to their own promoters *in vivo* (**FIG 5**) (9, 10).

**FIG 5:**
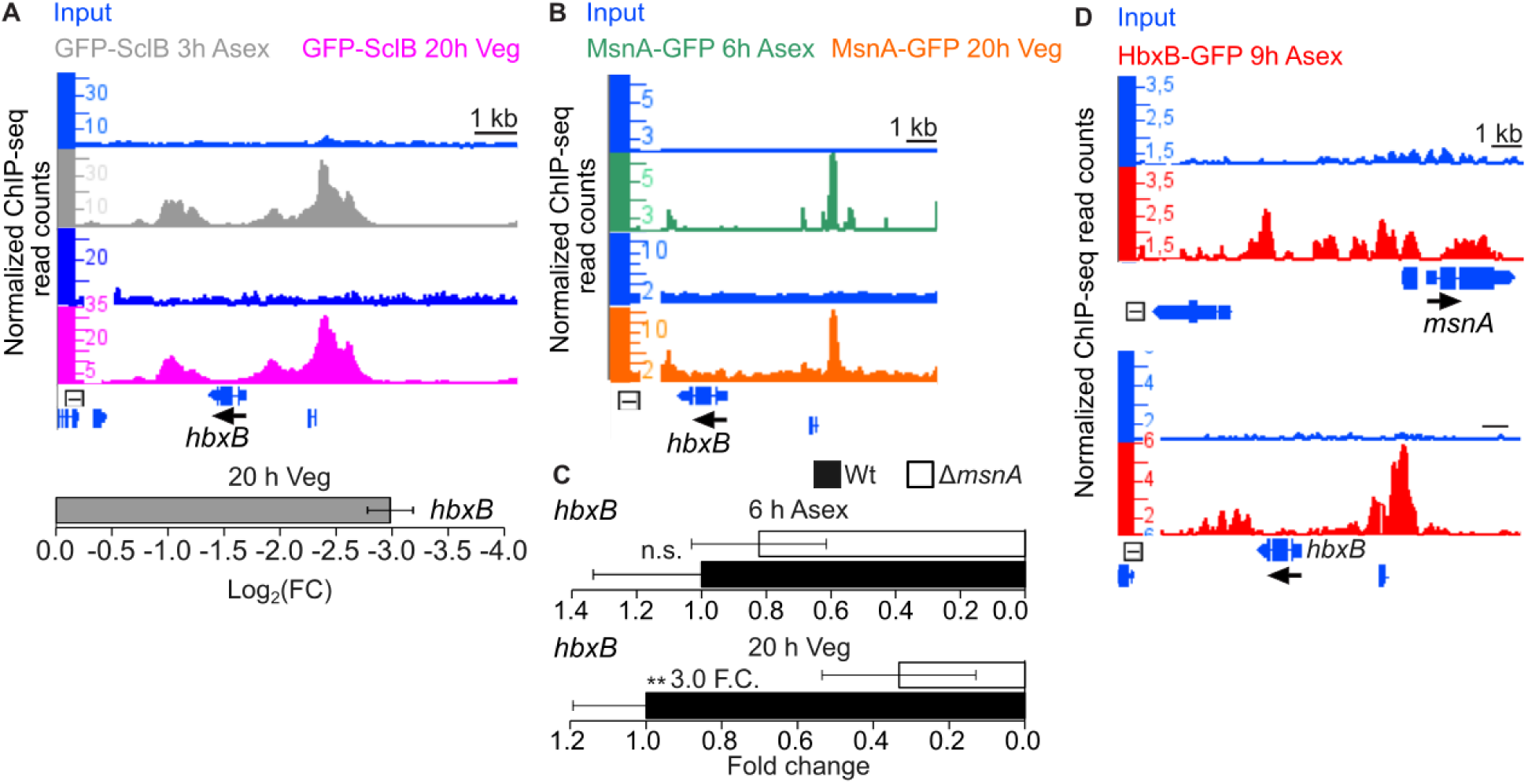
A transcriptional circuit between the HbxB, SclB and MsnA transcription factors, coordinates early steps in *A. nidulans* asexual development. (**A**) IGB views from ChIP-seq data with mycelia derived from the GFP-SclB strain grown either 20 h vegetatively (purple tracks) or 3 h under Asexual growth conditions (grey tracks), showing the *in vivo* association of SclB to the promoter of *hbxB* under both conditions. The ChIP-seq data used for this illustration came from analysis of the raw NGS sequencing data from Bastakis et al, 2025; deposited at NCBI under the BioProject ID: PRJNA1292721. Lower part of the panel shows the differential expression of *hbxB* as derived from the RNA-seq data of Δ*sclB* versus Wt under 3 h asexual growth published in the same study, Bastakis et al, 2025; deposited at NCBI under the BioProject ID: PRJNA1293339. (**B**) IGB views for ChIP-seq data with mycelia derived from MsnA-GFP strain after 20 h vegetative (orange tracks) or 6 h asexual growth (green tracks), showing the *in vivo* association of MsnA to the promoter of *hbxB* under both conditions. ChIP-seq data used for this panel derived from the analysis of the raw NGS sequencing data of the from Bastakis et al, 2025 PLoS Gen; deposited at NCBI under the BioProject ID PRJNA1260675. In all IGB views from the IGB the blue tracks represent the corresponding inputs (background signal), where no anti-GFP antibody was applied. Black arrows, showing the direction of the transcription of the gene above them and the black line represents a genome length of 1 kb. (**C**) Gene expression analysis by qRT-PCR, for the expression of *hbxB* with RNA derived from Wt and Δ*msnA* strains grown either 20 h under vegetative or 6 h under asexual growth conditions. The data depicted in the bar charts are averages and standard deviations of four biological replicates, each one of them consisting by four technical replicates. Statistical tests performed by Student’s t test and their interpretation is as: **p ≤ 0.01 and n.s.: not significant. (**D**) IGB views presenting ChIP-seq data for the *in vivo* binding of HbxB to its own promoter and to the promoter of *msnA*.

Developmental and stress responses in *A. nidulans* are mediated and processed by mechanisms that are controlling protein stability and signals transduction (34). Transcription factors constitute a central part of these mechanisms, as initially receivers of several signals and subsequently mediators of them. In that frame, the function of HbxB has been associated with several stress responses, particularly in *A. nidulans* asexual spores (18, 19). Hence, an analysis of the high-throughput data generated for this study was performed, with emphasis to stress-related genes. This uncovered a role of HbxB in either direct or indirect transcriptional control of genes coding for established stress associated proteins in hyphae growing under asexual growth conditions (**FIG S3**). This in not only in line with the role of HbxB in stress response and tolerance, but also enhances the previously mentioned molecular transcriptional circuit of HbxB, SclB and MsnA, since all of these TFs show responses in different kind of stresses such as for example oxidative stress (18, 19, 35–39).

Overall, the model that this study is proposing (**FIG 6**) presents HbxB as one of the main transcriptional regulators of *A. nidulans* asexual growth. HbxB has the ability to control the expression of genes, either directly via its *in vivo* association to their promoters or indirectly, in a genome-wide scale. Several of these genes encode essential proteins for asexual development such as SclB, PpoC, Flbs, VapA, MsnA and others. Furthermore, HbxB alongside with the MsnA and SclB transcription factors can be transcriptionally cross regulated. This trimeric relationship constitutes a new transcriptional circuit which potentially orchestrates several transcriptional responses for asexual growth. Additionally, HbxB exerts influence on the synthesis of various secondary metabolites, such as sterigmatocystin, austinol/dehydroaustinol and emericellamides during asexual growth. Lastly, HbxB is able to control the expression of stress-related genes, while the fungus is growing asexually. The findings of our study further enhance the mechanistic understanding of the transcriptional regulation of asexual growth. This knowledge can help in the future to develop better strategies against fungi that show detrimental effects to us or to species with great economical interest. It can further help to refine and develop better techniques for manipulating fungi with great biotechnological interest.

**FIG 6:**
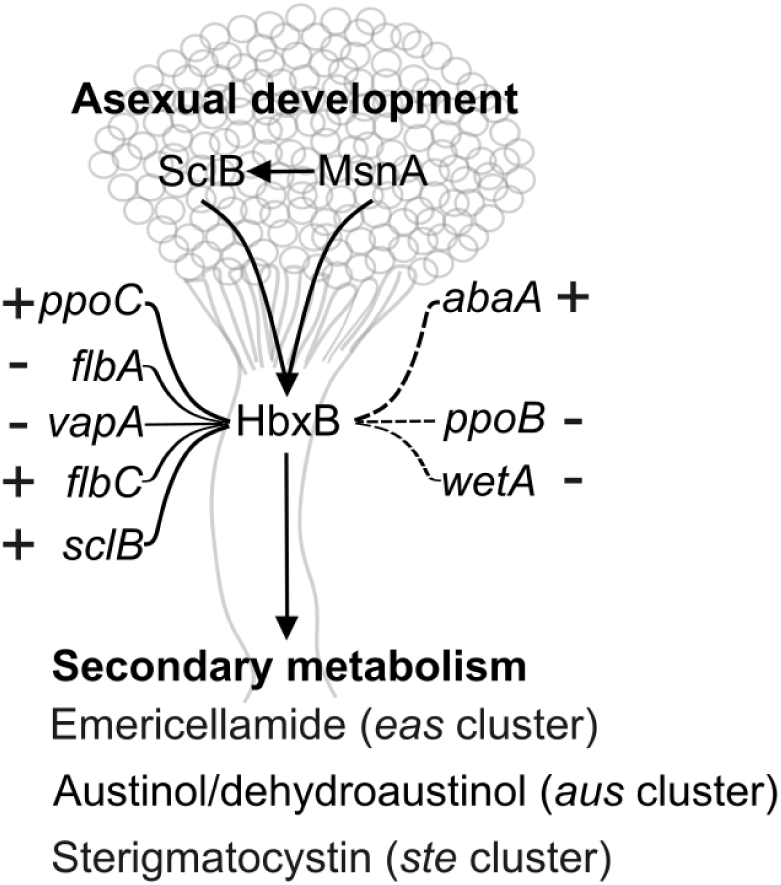
Transcriptional signals of asexual growth are initiated or mediated by the HbxB transcription factor. The graphical illustration depicts the prominent position that the HbxB regulator has during asexual development of *A. nidulans*. HbxB directly and indirectly influences transcription of genes during asexual growth, which code for established asexual regulators. Moreover, HbxB induces the synthesis of secondary metabolites with implications in development and defense of the fungus. Solid lines in the network illustrate direct *in vivo* transcriptional control. Punctuated lines in the network present indirect regulation by HbxB. The symbols (+) and (−) refer to induction and repression of the corresponding genes. The graphical model was done by employing the freely available vector-based software Inkscape (https://inkscape.org/).

## Material and Methods

### Growth conditions, medium and strains

*A. nidulans* strains used in this study were growing either under vegetative growth (spores inoculated liquid medium and followed shaking via on a rotary shaker for 20 h under light and 37 °C) or asexual favourable conditions (plates with *A. nidulans* spores let to grow upon solid medium for the indicated amount of time under light and 37 °C). The strains used for the ChIP-seq and the RNA-seq experiments were grown under identical growth conditions. In short mycelia from the corresponding strains growing vegetatively for 20 h and then they were transferred to solid medium plates for another 9 h under asexual growth conditions. After that it followed the collection of the mycelia and the corresponding downstream processing for each experiment. Any further modification to the growth conditions for the *A. nidulans* strains used in this study are mentioned directly to the corresponding parts and figure legends of the manuscript.

The composition of the minimal medium used to cultivate the *A. nidulans* strains was: 1% (w/v) glucose, 2 mM MgSO_4_, 1x AspA (7 mM KCl, 70 mM NaNO_3_, 11.2 mM KH_2_PO_4_, pH 5.5), 0.1% (v/v) trace element solution (76 µM ZnSO_4_, 178 µM H_3_BO_4_, 25 µM MnCl_2_, 18 µM FeSO_4_, 7.1 µM CoCl_2_, 6.4 µM CuSO_4_, 6.2 µM Na_2_MoO_4_, 174 µM EDTA) and pH 5.5 (40). When liquid medium was used, it was additionally supplemented with 0.1% (v/v) pyridoxine and 5 mM uridine. When solid medium was used, it was composed additionally with 2 % agar and 0.1% (w/v) uracil as well. The solid medium that was used after the transformation of *A. nidulans* protoplasts, was additionally containing 120 mg/mL nourseothricin, for the selection of the positive clones after the transformation. For the positive clones, selected after the transformation assay, there was a recycling of the selection marker, by growing the corresponding clones over a solid medium 0.5% (w/v) glucose and 0.5% (w/v) xylose correspondingly. The strain AGB551 (41) was used as the wildtype (Wt) for all the experiments in this study.

For the cloning purposes of this study it was used the DH5α strain of *Escherichia coli*. The cultivation of *E. coli* was in lysogeny broth (LB) medium (42) composed by 1 % tryptone, 0.5 % yeast extract, 1 % NaCl. There was an additional supplementation with 100 µg/mL ampicillin (final concentration) for the selection of desired positive clones after transformation with the preferred plasmid. Solid medium was additionally containing 2 % (w/v) agar. All strains of *A. nidulans* and *E. coli* used in this study are listed at the **S2 Table**.

### Plasmid manipulations and cloning

The whole processes of cloning (strategy, basic/backbone plasmids, kits, enzymes, sequencing, software etc) for the construction of the cassettes that were later used for transformation in *A. nidulans* strains, were performed as described in Bastakis et al., 2025 (9).

### Construction of the deletion cassette and the *ΔhbxB* deletion strain in *A. nidulans*

Amplification of the 5’- and 3’-flanking regions, was performed by using the MB252/253 and MB254/255 primer sets, correspondingly, using in both cases as template gDNA from Wt of *A. nidulans*. The amplicon for the 5’-flanking region yielded a fragment of 1781 bp and for the 3’-flanking region a fragment of 1524 bp. The 5’-flanking region was starting just after the translation start site and continued upstream towards the 5’-UTR and subsequently the promoter region of *hbxB*. The 3’-flanking region was starting just after the TGA, triplet encoding for the stop codon of *hbxB* and extended downstream from that point on. This cloning design targeted the complete deletion of the *hbxB* protein coding region after the transformation of the constructed cassette in to AGB551, Wt strain of *A. nidulans.* Both fragments were then cloned into the pME4696 backbone vector (43) producing the final plasmid of pMB37-1. It followed a verification of the proper sequencing for the cassette via Sanger sequencing. After that the cassette was excised with the *Mss*I (*Pme*l) restriction enzyme from the pMB37-1 and then was used for transformation in *A. nidulans* Wt strains. Finally, after the recycling of the selection marker, the whole process resulted to the AnMB21-1 A*. nidulans* strain.

### Assembly of the *hbxB::hinge::gfp* cassette and the *ΔhbxB* HbxB-GFP complementation strain in *A. nidulans*

The 5’-flanking region for this cassette was synthesized by two different fragments that where having compatibility introduced by proper primer designing and they been fused to one final piece performed via fusion PCR. The set of primers MB1337/MB1338 was used for the amplification of the first fragment of 1917 bp, starting just after the translation start site of the *hbxB* and expanding upstream covering the 5’-UTR and part of the promoter of the *hbxB* gene. As template for this amplification was used gDNA derived from the Wt, *A. nidulans* strain. The second amplicon of the 5’-flanking was amplified by using the primers MB793/MB794 to amplify the sequencing encoding for a linker/hinge (15 bp) followed by a sequence coding for the *gfp*. The overall size of the second amplicon was 729 bp. The plasmid pChS242 (44) was used as a template in the second amplification. The first and the second amplicon were then joined by fusion PCR using the set of the primers MB1337/MB794. The final fragment had a length of 2720 bp and it constitute the 5’-flanking region of the cassette. The 3’-flanking region was amplified using as template gDNA from the Wt strain of *A. nidulans* and the set of the primers MB1339/MB1340. The size of that amplicon was 2121 bp. The 5’- and 3’-flanking regions were finally clones into the pME4696 backbone plasmid producing the final vector of pMB38-1. After checking the sequence of corresponding cassette via Sanger sequencing, the cassette carried by this plasmid was excised by employing the *Mss*I (*Pme*l) restriction enzyme. It followed the transformation of the cassette into *A. nidulans* AnMB21-1 Δ*hbxb* strain. After the recycling of the selection marker carried by the cassette, the whole process led to the AnMB56-9 *A. nidulans* complementation strains.

### Transformations

The protocols used for the transformation in *E. coli* and in *A. nidulans* strains, were described by Meister et al., (43). Positive transformed clones of *A. nidulans* (those carrying the cassette in their genome), where further checked by Southern hybridization, performed as described by Southern 1975 (45). The labelling of the probes for the Southern hybridization was done by AlkPhos Direct Labelling Module (GE Healthcare Life Technologies, Little Chalfont, UK) according to manufacturer’s instruction.

### Phenotypical assays

To examine the development of the fungus under asexual growth conditions, 2000 for each strain were point inoculated in the middle of MM plates, supplemented with 0.1% (v/v) pyridoxine, 5 mM uridine and 0.1% (w/v) uracil. Subsequently, scans from the top and the bottom of the plates were made after the indicative number of days under the asexual growth.

To quantify the production of conidia for each strain under asexual growth conditions, initially the same number of spores from all strains were distributed equally on solid medium plates. It was used a minimum number of three biological replicates for each strain. Plates were growing under asexual growth conditions for the indicative number of days prior to the collection of the spores and the subsequent quantification by Coulter Z2 particle counter (Beckman Coulter GmbH, Krefeld, Germany).

### Secondary metabolites analysis

The preparation of the plates with the different strains of fungus was done as described by Liu et al. and left then to grow for 3-d under asexual growth conditions. Sampling, extraction of the secondary metabolites, running at the LC-MS/MS platform and the subsequent analysis were also performed as described by Liu et al., (46). The details of the detected secondary metabolites mentioned in the results are presented in **S5 Table**.

### DNA and RNA extraction

The isolation of the genomic DNA (gDNA) from mycelia growing overnight in liquid cultures at 37 °C and light, was performed according to Thieme et al., (35).

For the isolation of RNAs, 100 mL liquid medium was initially inoculated with 10^8^ spores for each strain; four biological replicates were used for each strain. Cultures then grew at the precise indicative conditions and mycelia at the end were collected and dried. A total of 100 mg dried mycelia for each replicate of each strain were snap frozen in liquid nitrogen. For the isolation of RNAs and the cDNA synthesis it was followed the procedure as described by Bastakis et al., (9).

### Quantitative real-time PCR

The gene expression of *hbxB* was assessed by employing a quantitative real-time polymerase chain reaction (qRT-PCR). RNAs derived from mycelia of Wt and Δ*msnA A. nidulans* strains either growing for 20 h under vegetative growth or initially growing under 20 h vegetative and then subsequently transferred for another 6 h under asexual growth conditions. The whole procedure (protocol, kits, instrumentation and analysis) that was followed is described by Bastakis et al., (9). The primers used in the qRT-PCR are listed at the **Table S1**.

### ChIP, sequencing and data analysis

For the ChIP experiment, initially, a total number of 5×10^8^ spores was inoculated in 500 mL liquid medium inside of 2 L flasks. Cultures were growing under constant shacking for 20 h under vegetative conditions. The produced mycelia were then transferred for additional 9 h on solid minimal medium plates under asexual growth conditions (light and at 37 °C). The complementation strains of Δ*hbxB* HbxB-GFP, expressing under its own promoter the *hbxB* fused with *gfp* (*green fluorescent protein*), was used for this experiment. Mycelia were promptly collected and fixed in solution containing 1 % formaldehyde for 20 minutes. The next steps of the ChIP protocol were performed as described in Sasse et al., (44) with the following modifications. Four independent biological replicates were used for the ChIP experiment. After the step of the shearing of chromatin each of the four samples were split in two parts. The first part was subjected to IP (immunoprecipitation) by applying the anti-GFP antibody (Abcam, ab290). The second part was not subjected to antibody, hence, served later as a negative control to subtract the background(noise) signal from each sample. However, both the IP and the input samples were treated in the same manner and pass through the same procedures till the end of the ChIP and the purification of the DNA. Reagents, instruments, kits and rest of the procedures for the ChIP experiment were identical as described in Sasse et al., (44).

The preparation of the ChIP-seq libraries and the following sequencing were performed according to Bastakis et al, (9) at the NGS-Integrative Genomics Core Unit (NIG), University Medical Center Göttingen.

The analysis of the produced ChIP-seq data was done by following the pipelines as presented in Sasse *et al*., 2023 (44) with some modifications that are following. Some parts of the analysis were completed by the GALAXY platform (47) as provided by the GWDG (Gesellschaft für wissenschaftliche Datenverarbeitung mbH Göttingen). Mapping of the raw sequences data to the *A. nidulans* genome (downloaded from fungidb.org: FungiDB-46_AnidulansFGSCA4_Genome.fasta) was performed by Bowtie2 (48). For the subsequent detection of the statistically significant peaks among each of the four independent sets of analysis was used the MACS2 tool (49). For the generation of files compatible for visualization of the ChIP-seq peaks to the Integrative Genome Browser (IGB) (50) was used the bamCoverage tool from the deepTools2 package (51). The analysis of the distribution of the statistically significant peaks over different genomic elements was performed by the R-based tool ChIPseeker (52). The *de novo* motif discovery was performed as described in Sasse et al., (44) by the RSAT webtool (24).The GO-enrichment analysis was performed with the ShinyGo v0.82 webtool (53). The Venn diagrams were made by using the InteractiVenn web tool (54). The raw sequencing data for the ChIP-seq have been deposited at the SRA archive of NCBI [BioProject ID: PRJNA1330218].

### RNA-seq and data analysis

RNA-seq libraries and the sequencing, were performed at the NGS-Integrative Genomics Core Unit (NIG), University Medical Center Göttingen. The construction of the libraries it was followed the pipeline by Szemes et al., (55). The subsequent RNA-seq analysis was performed in GALAXY platform and in R as provided by GWDG. Raw sequencing reads were mapped in the *A. nidulans* genome (downloaded from fungidb.org: FungiDB-46_AnidulansFGSCA4_Genome.fasta) by using the Bowtie2 (48). Next, were prepared matrices for the subsequent part of the RNA-seq analysis, by using the htseq-count tool (56). These matrices were then used by DESeq2 (57) to calculate the expression of the DEGs between the Δ*sclB* versus the Wt samples. The heatmaps presented in this study were generated by the R-based package complexHeatmaps (58). The raw sequencing data for the RNA-seq have been deposited at the SRA archive of NCBI [BioProject ID: PRJNA1330713].

### Microscopy

Fluorescence microscopy was performed as described in Bastakis et al., (9). The nuclear localization of the HbxB-GFP was assessed by the by the global nuclear marker of H2A tagged with the RFP (RFP-H2A). The Wt strain (negative control), was also expressing RFP-H2A. The cassette for the ectopical expression of this marker (*^p^gpdA::intron::mrfp::h2A*(*cDNA*) was excised from the plasmid pME3173 (59). Through transformation in the corresponding *A. nidulans* strains occurred the integration of the gene encoding the nuclear marker to the genome resulting into the strains, AGB1718 (10) and AnMB57-1.

## Figures processing

All figures were processed by the vector-graphics editor Inscape (Inkscape Project, 2020; Inkscape, available at https://inkscape.org).

**S1 Table: Primers used in this study.**

**S2 Table: Strains (*E. coli* and *A. nidulans*) used in this study.**

**S3 Table: Genes where HbxB found to be associated reproducibly (*in vivo*) to their promoters in the ChIP-seq experiment.**

**S4 Table: Differentially expressed genes as found in the analysis of the RNA-seq experiment performed with RNAs derived from Δ*hbxB* and Wt strains growing for 9 h under asexual growth conditions.**

**S5 Table: Detected Secondary metabolites.**

**S6 Table: List of 44 stress related genes in *A*. *niduans*.**

## Acknowledgments

We thank Dr. Gabriela Salinas and Fabian Ludewig from the NGS-Integrative Genomics Core Unit (NIG), University Medical Center Göttingen for their excellent support on our NGS-based approaches. We would also like to thank Gabriele Heinrich, Vincent Anthony Czarnowski Corona and Victor Flesch for their technical support.

## Supplementary material

**FIG S1:**
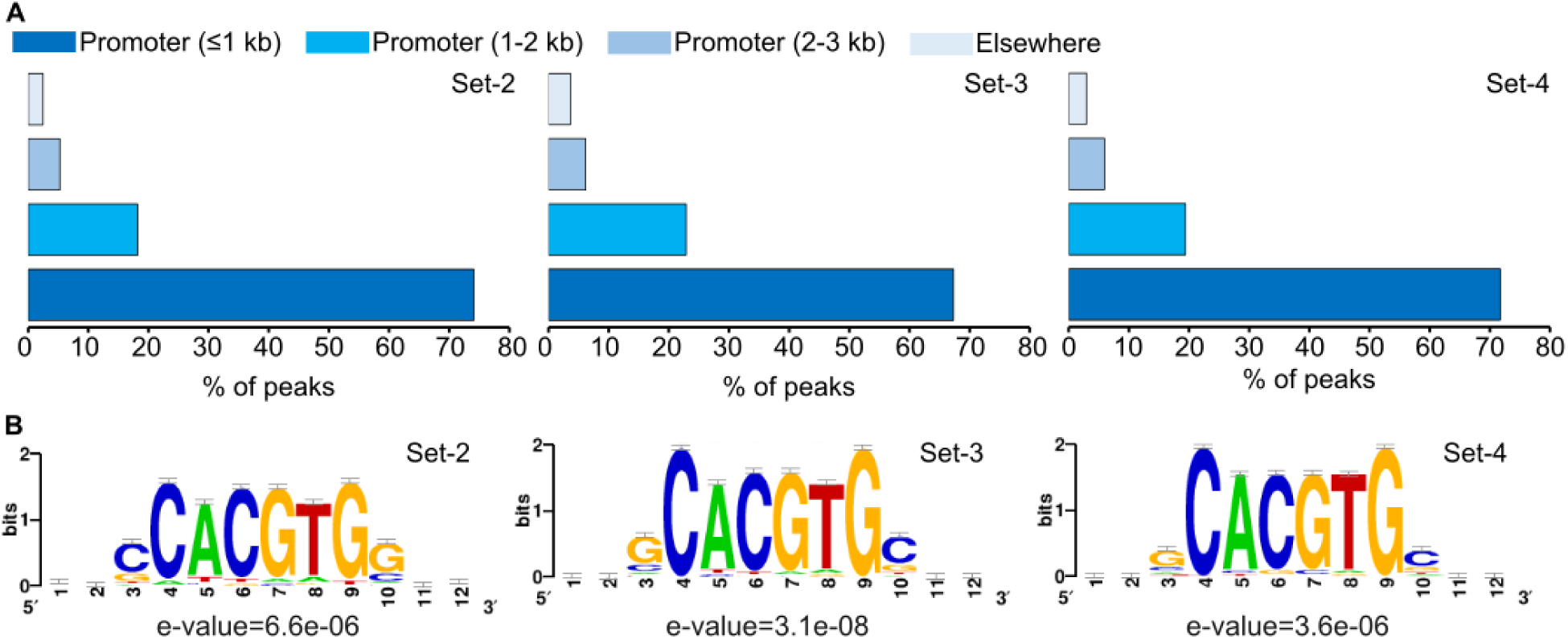
HbxB predominantly associated with promoter regions spanning up to 1 kb from the TSS, via recognizing a specific DNA element. (**A**) Bar charts illustrating the distribution of the ChIP-seq data, from set-2 till set-4, over different genetic elements of the *A. nidulans* genome. (**B**) Motifs presenting the DNA element recognized by HbxB with which it is associated *in vivo*. The *de novo* motif discovery for the identification of this motif was performed with the RSAT-web tool, using as input 100 bp sequences lying underneath the summits of the top 150 scored peaks (binding positions of the HbxB) for each independent set of analysis (set-2 till set-3).

**FIG S2:**
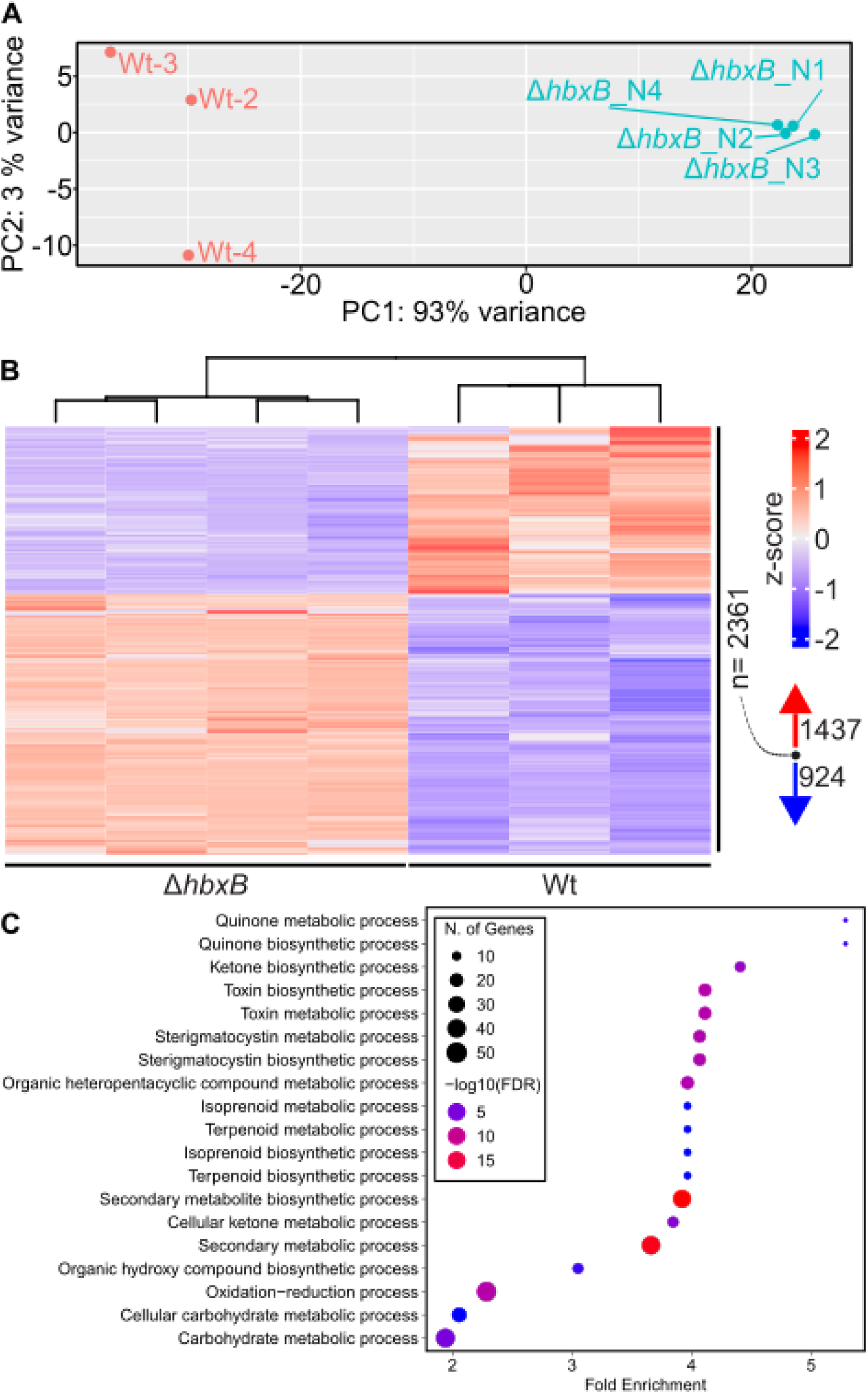
HbxB influences the expression of 2361 genes from the *A. nidulans* genome during asexual growth of the fungus. (**A**) Principle component analysis reveals a distinct clustering among the samples from RNA-seq that derived from Wt and Δ*hbxB* strains grown for 9 h under asexual growth conditions. (**B**) Heatmap illustrating the 2361 genes found to be differentially expressed (applied cut offs: p < 0.05 and −1 ≤ log_2_FC(Fold Change) ≥ 1) in the RNA-seq performed with RNAs derived from mycelia of Δ*hbxB* and Wt strains, grown under asexual conditions for 9 h. (**C**) gene ontology(GO)-enrichment analysis, performed by the ShinnyGO web tool, in terms of biological processes (BP), for the 2361 DEGs of the RNA-seq. Black circles of different size, present the size in a number of the genes that belong to each category in all the identified overrepresented groups. The blue-red gradient-colored circles depict the statistically significance that each identified category has (−log10(FDR)).

**FIG S3:**
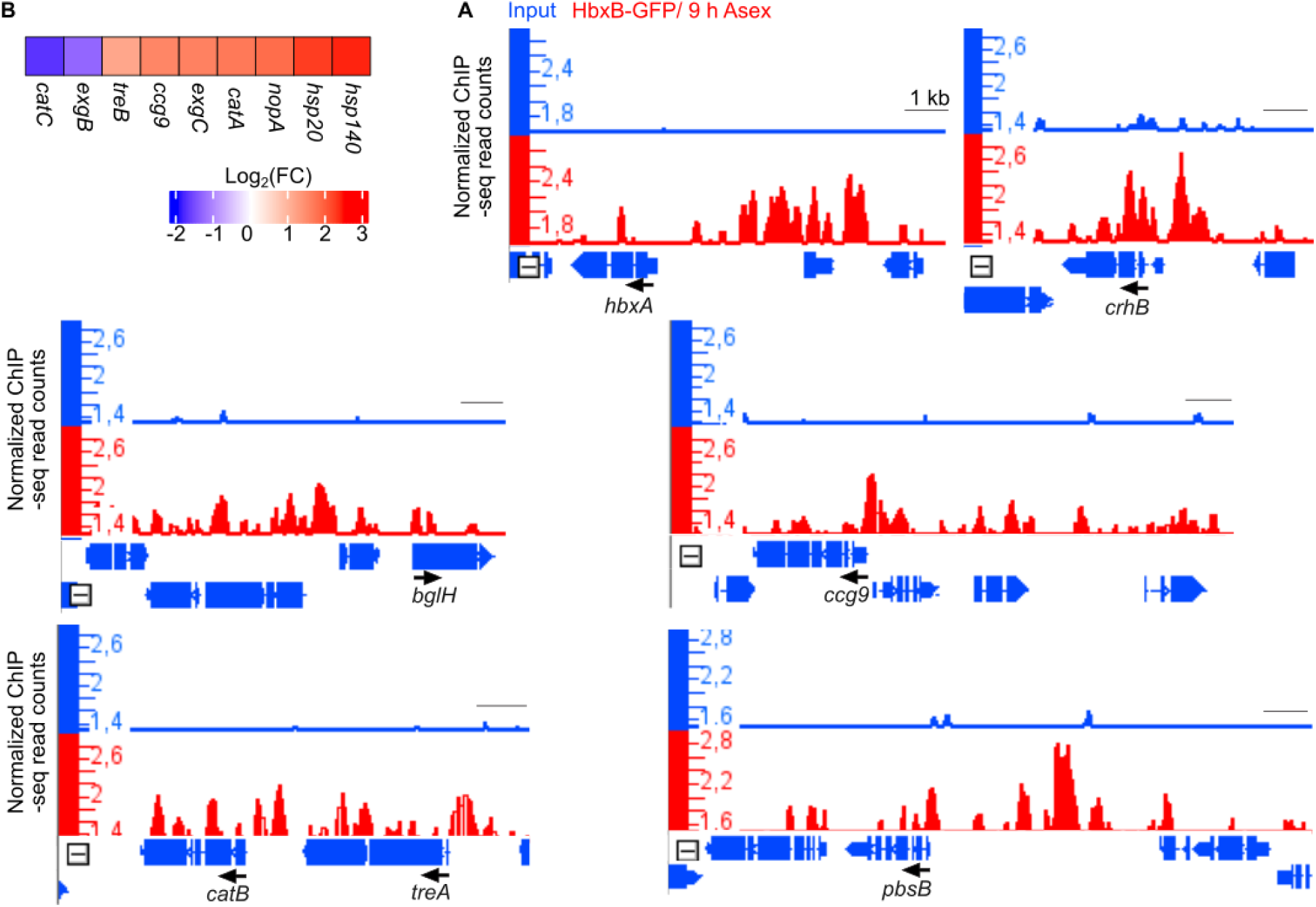
HbxB influences the expression of genes encoding several stress-related proteins in *A. nidulans* during the asexual growth. (**A**) Heatmap showing the differential expression of genes coding for stress-related proteins, from the RNA-seq that was performed with RNAs derived from Δ*hbxB* and Wt strains grown for 9 h under asexual conditions. For the analysis were applied cut offs: p < 0.05 and −1 ≤ log_2_FC(Fold Change) ≥ 1). (**B**) Screen shots from IGB illustrating one representative independent set of analysis among (out of four in total) the ChIP-seq data for the *in vivo* binding of HbxB to promoters of stress-related genes. Red tracks showing the mapped reads of the ChIP-seq data derived from the IP of HbxB-GFP with the anti-GFP antibody. Blue tracks are presenting the corresponding input of this sample, where no anti-GFP was added to (background signal). Black arrows indicate the direction of the transcription of the gene above it and black line represents a genome length of 1 kb.

**FIG S4:**
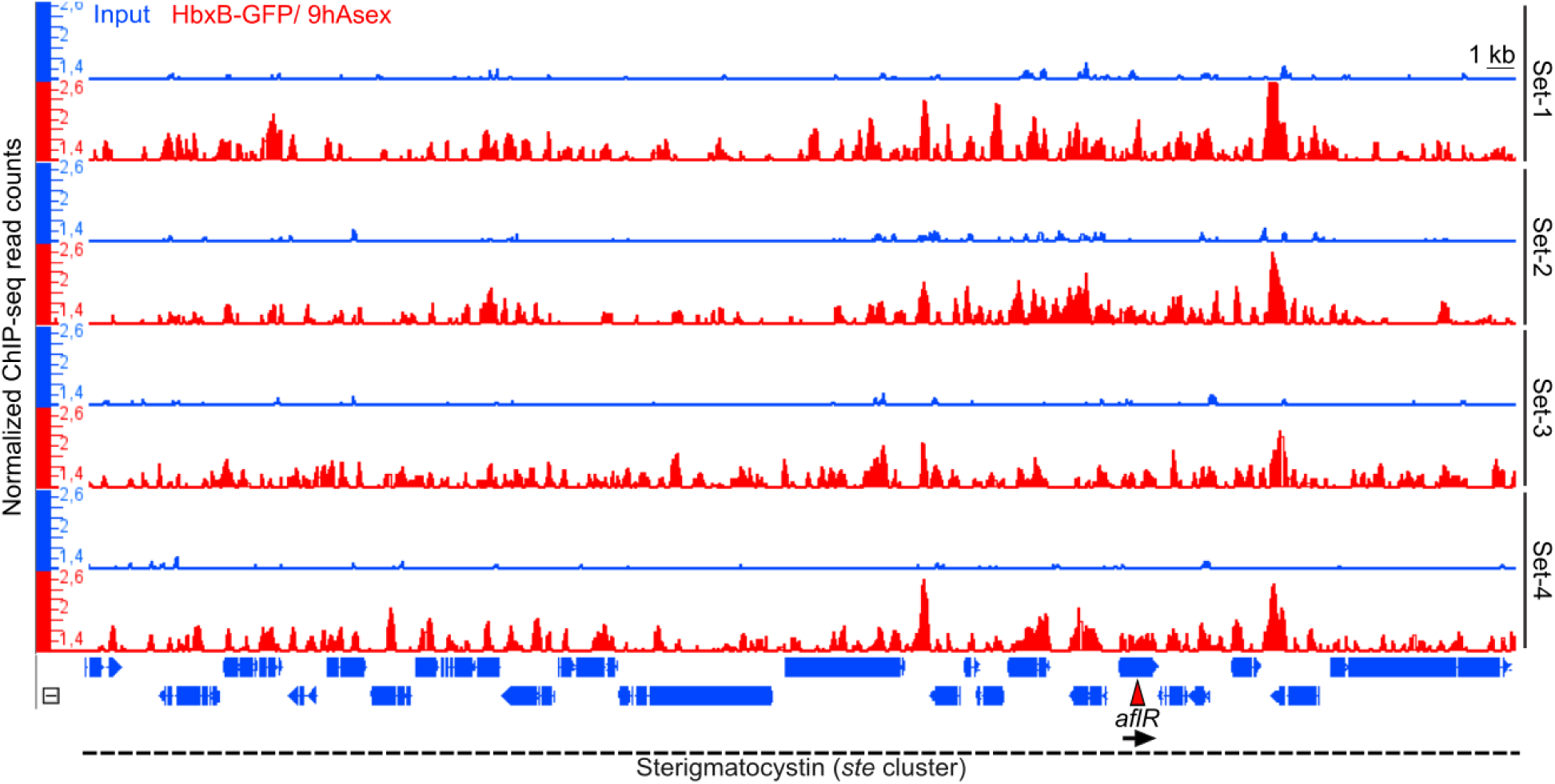
Sterigmatocystin (*ste*) gene cluster displays multiple binding occurrences of the HbxB transcription factor. IGB views, showing the *in vivo* binding of HbxB to the promoter regions of several gene members of the cluster. Highlighted by a red triangle is the *aflR* gene encoding the main transcriptional regulators of the whole cluster. The picture shows that only in three, out of four in total, independent sets of the ChIP-seq analysis, there were statically significant (p<0.05) peaks for the *aflR* gene identified. Red tracks showing the mapped reads of the ChIP-seq data from the IPs of HbxB-GFP with the anti-GFP antibody; blue tracks illustrating the corresponding inputs of these samples, where no anti-GFP antibody was added (background signal). Black arrow is showing the direction of the transcription of *aflR* and black line representing a genome length of 1 kb of the genome.

